# Survivability and Life Support in Sealed Mini-Ecosystems with Simulated Planetary Soils

**DOI:** 10.1101/2023.11.02.565408

**Authors:** Tsubasa Sato, Ko Abe, Jun Koseki, Mayumi Seto, Jun Yokoyama, Tomohiro Akashi, Masahiro Terada, Kohmei Kadowaki, Satoshi Yoshida, Yosuke Alexandre Yamashiki, Teppei Shimamura

## Abstract

Establishing a sustainable life-support system for space exploration is challenging due to the vast distances, costs, and differing environments from Earth. Using insights from the Biosphere 2 experiment, we introduced the “Ecosphere” and “Biosealed” systems in custom containers to replicate Earth’s ecosystems, suggesting feasible space migration through transplanting Earth-like biomes.

Over four years, we gained deeper insights into these enclosed ecosystems. Moisture deficiency was a major obstacle to plant growth, which we addressed by incorporating a groundwater layer in the containers. We underscored the critical role of microorganisms in building and sustaining these ecosystems. However, temperature spikes from sunlight threatened stability. Our experiments confirmed fruit flies’ survival on plant-produced oxygen and photosynthetic bacteria. Interactions between plants, microbes, and simulated space soils were examined. Detailed analysis unveiled diverse microbes shaping both confined and simulated space environments. Major findings include the symbiotic relationship of plants with cyanobacteria, the potential of LED lighting in sun-limited missions, and challenges with ethylene gas and moisture. Microbial integration in rough soils holds promise for seed germination, but understanding their role in space soils is crucial.

Our research offers a comprehensive foundation for future space life-support systems and underlines potential concerns about microbes affecting human health.

## Introduction

Establishing a life-sustaining system for extraterrestrial planets is a significant challenge in humanity’s quest for space exploration. Within the vastness of space, where sending supplies from Earth becomes a daunting economic task, especially for spacecrafts and space colonies, there’s a pressing demand for technologies that allow self-sufficiency in terms of food and oxygen. In addition, the alien environments of these extraterrestrial planets differ starkly from Earth, characterized by intense radiation, distinct atmospheres, and a stark scarcity of water resources. Therefore, there is an urgent need for research on how we harness the finite cosmic resources in order to develop and maintain sustainable, recycling-oriented ecosystems (Fig. 1a).

**Figure 1.**
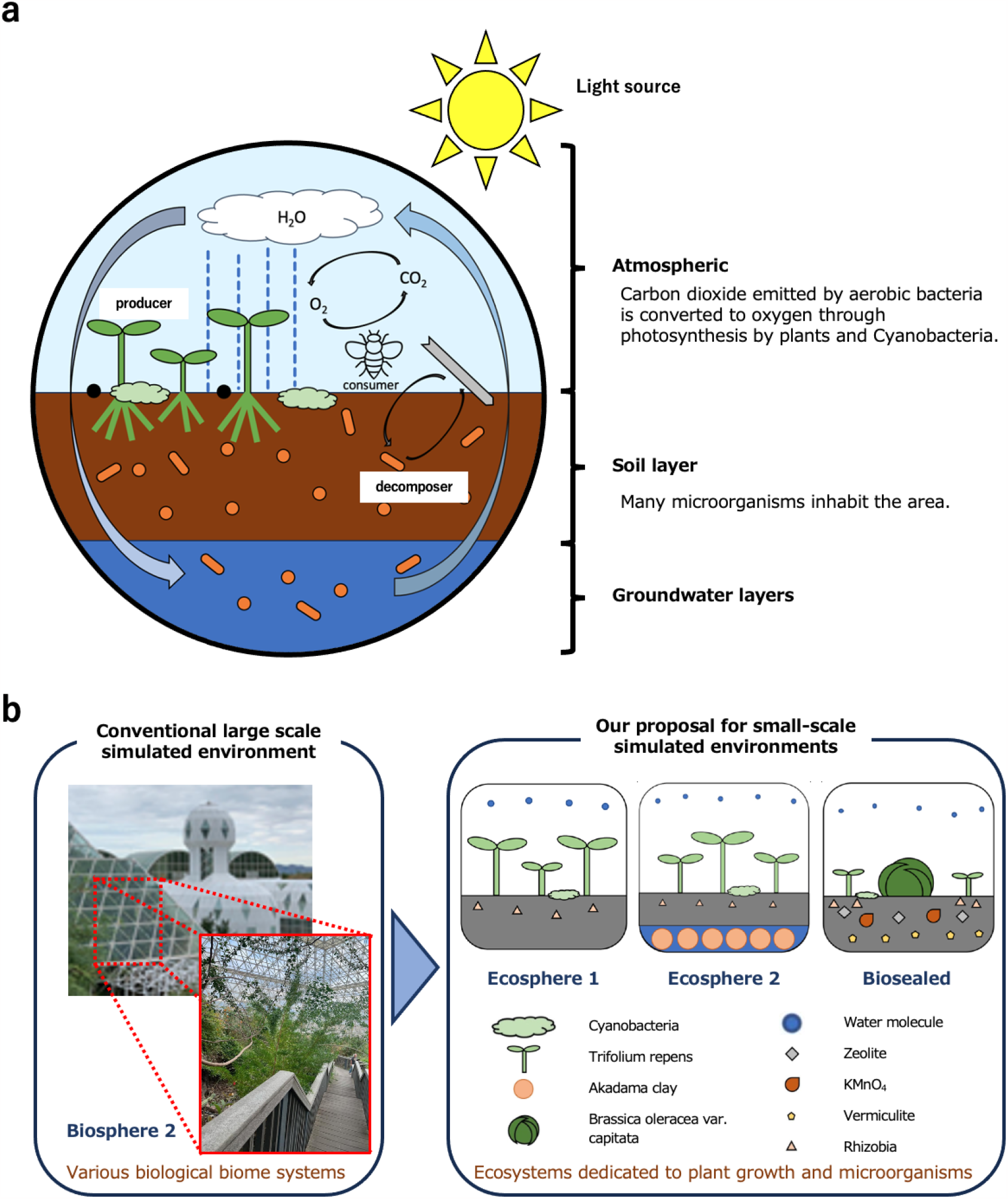
Concept of This Study. (a) Conceptual diagram of the research. (b) Introduction of the Ecosphere and Biosealed created in this study, along with a mention of the prior study, Biosphere 2.

A pioneering effort to construct and maintain such sustainable ecosystems in enclosed spaces is epitomized by Biosphere 2 (Allen, 1993, Fig. 1b, left panel, Allen, J. Biosphere 2, Nelson, M. 2021, Allen JP et al., 2003), located in Oracle, Arizona, USA. This facility served as an experimental ground to simulate potential ecosystems where humans might inhabit other celestial bodies and as a research site for studying changes in Earth’s ecosystems. Numerous closed-environment experiments were conducted here, addressing the sustainability of ecosystems and the psychological and physical well-being of humans, yielding invaluable data.

However, due to structural flaws leading to changes in soil microbial communities, oxygen levels dropped, adversely affecting many aerobic organisms. Food shortages and interpersonal tensions among researchers also arose, leading to the termination of the experiment after two years. These experimental outcomes underscored the need for a reduced ecosystem transfer concept for sustainable space colonization. They also highlighted the many challenges in maintaining a long-term sustainable ecosystem.

In this study, we introduced and designed compact, natural, closed-loop ecosystems called “Ecosphere” and “Biosealed” to address the challenges associated with sustaining life in enclosed spaces. Our main experiments focused on plant growth and survival, the presence and role of microorganisms, and cultivation using simulated extraterrestrial soil. Through these experiments, we aimed to gain a better understanding of the mechanisms and challenges of sustaining life in a closed environment and to identify potential solutions (Fig. 1b, right panel). Furthermore, growing plants in extraterrestrial environments, especially on distant planets and moons, is a crucial step in human space colonization (Meeboon et al., 2022, Fujiwara et al., 2013). Concurrently, we delved into plant cultivation in simulated extraterrestrial soils to explore the presence and impact of microorganisms (Byon et al., 2022, Macedo et al., 2009, Bernsten et al., 2007, Nishiwaki et al., 2022).

### Natural Circulation in a Sealed Space by Ecosphere 1

Inspired by Biosphere 2, we developed a sealed container named “Ecosphere 1”, with enhanced airtightness. This container is made of glass and is sealed with melted rubber or silicone for the lid. It is designed to be more compact than its predecessors, aiming to identify challenges in a basic closed ecological system by emulating a natural environment.

To investigate the characteristics of natural circulation in the closed space of Ecosphere 1, we enclosed nutrient-rich soil collected from a natural environment, and seeds of clover, a leguminous plant with nodules, into the Ecosphere 1. As a result, mainly clover growth was observed. Emphasizing the role of microbes in maintaining a stable natural ecosystem, experiments were conducted using soil containing naturally derived microbes. Finally, Ecosphere 1 was placed outdoors on a wooden board at a height of about 1 meter above the ground to avoid heat from asphalt and concrete. The natural cycles inside it were then observed for four years, reflecting the seasonal changes in Japan.

Our analysis revealed the coexistence of plants and cyanobacteria, establishing diurnal oxygen supply and natural circulation through their activity. We also observed the decomposition process of withered clover stems and leaves. Mass conservation tests further confirmed the absence of flaws in the Ecosphere 1 structure (Sup. Fig. 1a,b, Sup. Tab. 1,2). These results suggest the viability of sustaining a unique ecosystem in a confined space and underscore the integrity of the Ecosphere 1 design.

Remarkably, Ecosphere 1 adapted to the diverse external environment of Japan’s four seasons, exhibiting its own natural cycles (Fig. 2a). In the first spring, only soil, water, and seeds were introduced into Ecosphere 1, and it was placed in a sunny location. Within 2-3 days, the clover seeds germinated. As summer approached and internal temperatures rose, most plants inhabiting the surface perished due to the heat retention of the sealed glass container. However, decomposers breaking down the dead plants were identified (Sup Fig. 2a). Furthermore, Cyanobacteria proliferated, covering both the surface and the ground (Fig. 2b). Their photosynthesis likely provided oxygen, possibly sustaining aerobic bacteria (Sup Fig. 2 b,c). In autumn, new shoots emerged from presumed rhizomes beneath the soil, while Cyanobacteria receded into the earth (Fig. 2c). Though winter witnessed some plants perishing at the end of their life cycles, others overwintered. Cyanobacteria flourished once more, enveloping the container surface (Fig. 2d). By the following spring, it became evident that new shoots sprouted from seeds left behind by the previous generation (Fig. 2e).

**Figure 2.**
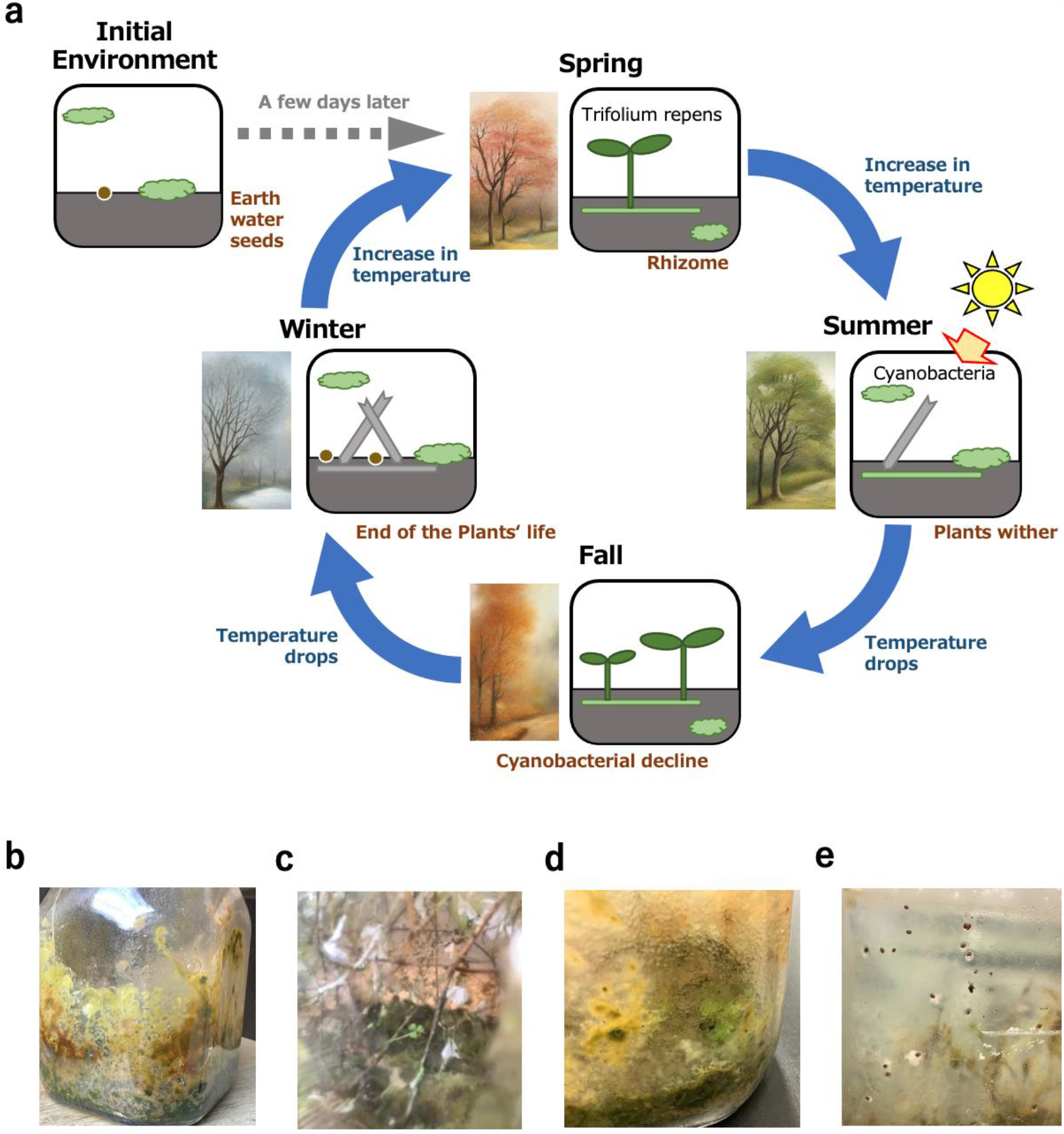
Natural Cycle Observed in Ecosphere 1 Under Natural Conditions. (a) Conceptual representation of the natural cycle within Ecosphere 1. (b) Observations of cyanobacteria proliferation during summer. (c) Appearance of regrowing clover in autumn. (d) Increase in cyanobacteria during winter. (e) Residual clover seeds.

### Growth Rate of Plants in a Sealed Environment

To closely examine the influence of a sealed environment on the growth rate of plants, observations were made within the Ecosphere 1 from January 13, 2021, to April 25, 2021, spanning approximately 15 weeks (Fig. 3a). In the sealed space, plants grew only about 10 cm over 15 weeks, whereas in an open environment, they achieved this growth in merely 5 weeks. This stunted growth is believed to be due to the accumulation of the plant hormone ethylene and a lack of essential moisture for plant growth (Drenovsky et al., 2004). It is speculated that the buildup of the plant hormone in the closed environment adversely impacted plant development. Additionally, given the characteristics of the Ecosphere, the total water content remained constant. As plants grew, the soil moisture content diminished, which was thought to further inhibit growth.

**Figure 3.**
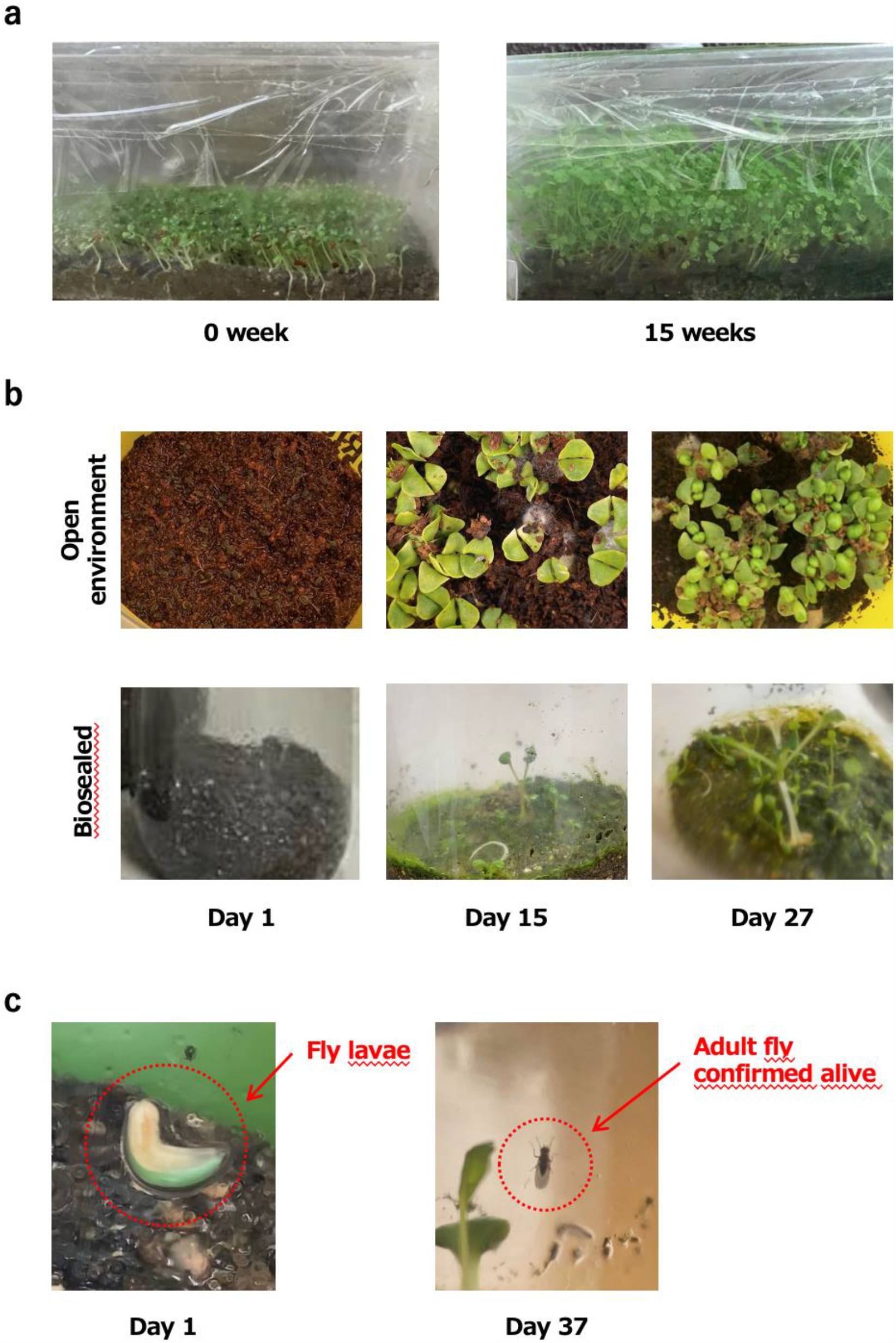
Results of Plant Growth Experiments in Sealed Environments & Survival of Yellow Drosophila Fly. (a) Growth of clover in a sealed environment. (b) Plant growth under LED light conditions. The top image showcases basil growth in an open environment with LED lighting, while the bottom shows cabbage growth in a sealed environment with LED lighting. (c) Representation indicating the sustainability of the yellow Drosophila fly in a sealed environment. The left image is from the first day where larvae were visible, while the right image is from the last observed moment, showing them flying within the container.

Drawing inspiration from these insights, we revamped Ecosphere 1 and birthed “Ecosphere 2”, endowed with an expansive underground aquifer (Sup. Fig. 3). Thanks to this new structure, the soil within Ecosphere 2 remained consistently moist regardless of seasonal changes or the growth status of the plants. Within Ecosphere 2, while plants stretched taller, their leaf dimensions remained steadfast, possibly under the influence of the ethylene hormone.

### Plant Cultivation in a Sealed Environment Using LED Lighting

During outdoor maintenance of Ecosphere1, many plants succumbed to the intense summer heat. In extraterrestrial environments, such as the moon, there is no thick atmosphere like on Earth. Consequently, the heat from sunlight is directly transferred, compounded by intense radiation. In response, we experimented with plant cultivation using the SP312 LED light (NARRNA), which has less heat impact than natural sunlight. We chose basil (*Ocimum basilicum*) and cabbage (*Brassica oleracea var*.*capitata*) for the study, given their resistance to diseases and pests, and their apt sunlight requirements and cold tolerance.

Initially, to verify the suitability of the LED lights for plant growth, we conducted germination and growth experiments with basil in open conditions, and with cabbage in enclosed environments (Fig. 3b, top panel). By day 15, the cabbage germinated normally, and the basil growth was progressing well (Fig. 3b, bottom panel). An observation of cabbage germination in an open environment, without soil, confirmed its normal germination by day 15 (Sup. Fig. 4).

Building on these results, we attempted plant cultivation in a closed space. We planted clover and cabbage simultaneously, monitoring their growth under the LED lights. Both plants were able to sustain life. However, a characteristic challenge in enclosed spaces was observed: the tendency for leaves to be smaller. Nevertheless, this tendency was also observed in natural sunlight conditions, leading us to conclude that it wouldn’t pose issues for subsequent research.

### Observations on Sustaining Life in Sealed Environments

In the preliminary stages of our experiment with Ecosphere1, a dead fly was discovered one month after the experiment’s inception. However, no other fly remains were detected thereafter. Based on this initial test, we introduced an advanced model, “Biosealed”, incorporating clover, cabbage, and artificially added Cyanobacteria. Biosealed, made of glass, doesn’t feature an underground aquifer; instead, it uses soil enriched with vermiculite to enhance moisture retention.

To this Biosealed model, we introduced eggs of the fruit fly (Drosophila melanogaster), assessing life sustenance potential under conditions similar to the previous experiment. Thanks to a consistently maintained temperature in a stable environment, larvae were identified within a week. Under these conditions, the fruit fly eggs hatched and matured into adults. Notably, the last observation of a living fruit fly was on day 37 post-introduction. Considering that the average lifespan of a fruit fly in natural conditions is known to be 28 days, we can infer that life sustenance in the enclosed environment was a success (Fig. 3c). We speculate that the extended life span could be attributed to the more stable conditions than those found in nature.

From the above findings, it is evident that life can be sustained in enclosed environments for a definite period.

### Plant Cultivation in Exoplanetary Simulated Soils

In this study, we examined the variations in plant cultivation environments within sealed spaces using multiple extraterrestrial simulated soils. Notably, in the microbe-free Ryugu simulated soil, germination of plants was not observed (Fig. 4a). Given that the aqueous pH value of the relevant simulated soil stood at 8.45, we ruled out pH levels as the inhibiting factor for germination. Observations of darkened seeds and the presence of fine particles in the Ryugu simulated soil led us to postulate that these particles might have adhered to the clover seeds, potentially inhibiting their respiration.

**Figure 4.**
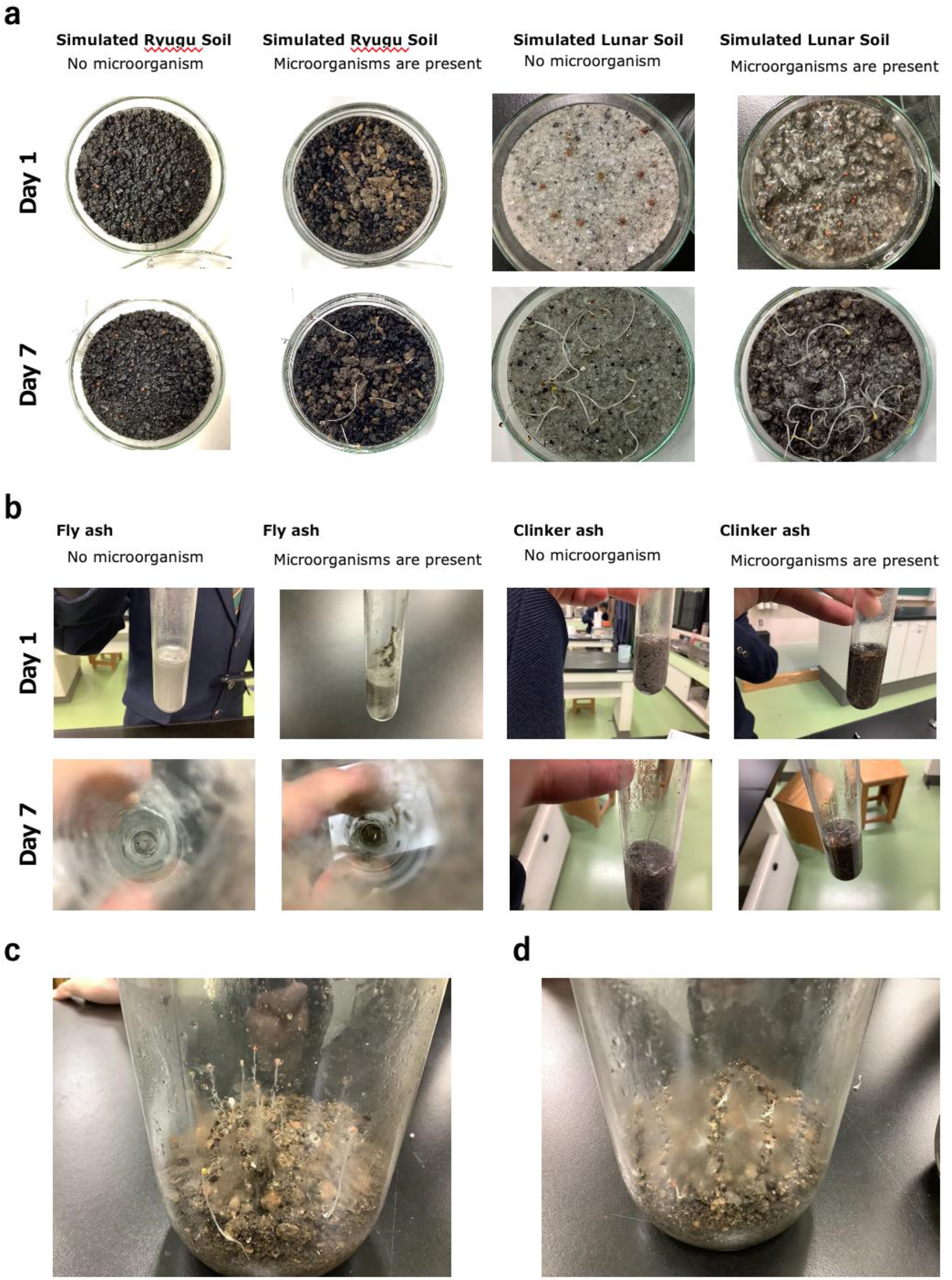
Germination Experiments of Plants in Exoplanetary Simulated Soils & Importance of Microbial Presence. (a) Germination rate comparison between plants in exoplanetary simulated soils with similar components, with and without introduced microbes. (b) Plant cultivation experiments under conditions resembling lunar soil, using Fly ash and Clinker ash. (c) Plant growth in a sealed environment without sterilizing the microbes. (d) Plant growth in a sealed environment post microbial sterilization.

Contrastingly, in the microbe-introduced Ryugu simulated soil, 6 out of 10 seeds successfully germinated (Fig. 4a). This suggests the potential of microbes to ameliorate the extreme conditions of the Ryugu simulated soil, laying the groundwork for further analysis. In the lunar simulated soil, irrespective of the presence of microbes, all seeds germinated. These results have been consolidated in Fig. 4a.

In both Fly ash and Clinker ash, germination and growth of clover were confirmed regardless of microbial presence (Fig. 4b). However, Fly ash, owing to its fine particles, exhibited heightened water repellency, posing a constraint to plant growth. Building upon prior research which reported difficulties in microbe fixation to soil (Paul et al., 2022), efforts to increase the porosity of Fly ash and Clinker ash were undertaken. Initially, Fly ash mixed with water was found to be fragile and had issues with excessive water absorption, rendering it unsuitable (Sup Fig. 5e). In contrast, solids made from cement and bentonite were porous but lacked adequate moisture retention, showing no signs of plant germination. Using a bonding agent, Booncrete exhibited moderate water absorption (Sup. Fig. 5h,j,k), yet its excessive hardness was deemed unsuitable for plant cultivation, indicating room for improvement.

**Figure 5.**
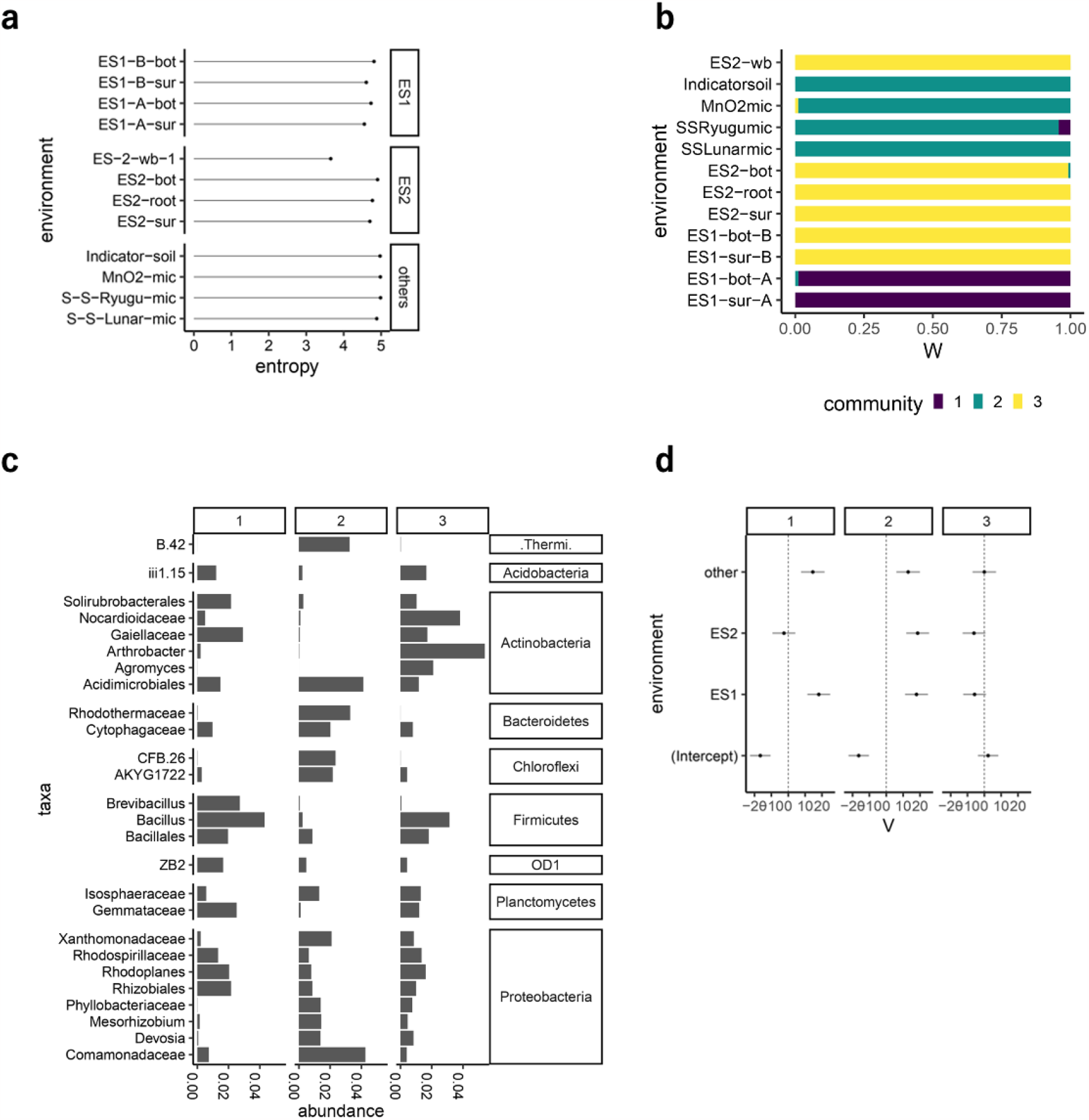
Metagenomic Analysis Results to Examine the Impact of Microbes on Plant Growth. (a) Entropy for each environment. (b) Microbial community composition for each environment, as estimated by BALSAMICO. (c) Predominant microbes in each community, as inferred by BALSAMICO. (d) Effects of environmental factors on the estimated microbial community composition by BALSAMICO (expressed as log odds ratio).

To evaluate the extent to which microbes contribute to forming a plant cultivation environment in sealed spaces, a comparative experiment between sterilized and non-sterilized soils was conducted. Consequently, non-sterilized soils showed favorable growth of clover (Fig. 4c), whereas growth in sterilized soils was minimal (Fig. 4d). These findings underscore the significance of microbial presence in improving conditions within extreme and sealed environments.

### Plant Growth and Microbial Interactions in Sealed Environments

To elucidate the interactions between plant growth and microbes, we conducted a metagenomic analysis. Initially, the relative abundance of microbes was depicted in Figure 5a using entropy (alternatively known as ASV-based Shannon diversity), a measure commonly utilized to indicate biological diversity and distribution. Ecosphere samples (ES1, ES2) displayed lower entropy compared to other samples, suggesting a pronounced presence of specific bacteria within the ecosphere. However, the time elapsed post-microbial introduction into the Indicator soil’s simulated soil was shorter than for ES1 and ES2, suggesting we might be observing a transitional state of microbial communities. Indicator soil is primarily composed of heterotrophic microbes. In the case of endogenous heterotrophic succession, it’s proposed that an intermediate supply of all limiting resources and a peak diversity in the organic carbon pool may occur (Fierer et al., 2010).

We employed BALSAMICO (BAyesian Latent Semantic Analysis of MIcrobial COmmunities) (Abe *et al*., 2021) to identify bacterial communities from microbiome data and to explore the association between environmental factors and these communities. BALSAMICO integrates a non-negative matrix factorization approach, factoring in environmental influences to model community structures. As illustrated in Fig. 5b, the analysis reveals that while bacterial community 1 predominantly characterizes ES1, ES2 is mainly defined by bacterial community 2, and the simulated soil is dominated by bacterial community 3. Intriguingly, the significant difference in bacterial communities between ES1 and ES2 suggests that humidity might have a profound impact on the bacterial community composition. Water activity is a known determinant of microbial communities (Drenovsky et al., 2004; George et al., 2021), which might play a more significant role than other environmental factors in distinguishing ES1 and ES2.

Delving deeper into the bacterial communities, Community 1, as evident from data analysis in Fig. 4c,d and Sup. Tab. 2, is primarily dominated by aerobic chemoautotrophic bacteria such as *Brevibacillus* from the phylum *Firmicutes*. Notably, the presence of oxygenic photosynthetic nitrogen-fixing bacteria, like *Leptolyngbya* from the phylum *Cyanobacteria*, is suggested. In Community 1, not only plants but also these bacteria might play a role in supplying both oxygen and organic compounds within the ecosphere. In contrast, Communities 2 and 3 have fewer oxygenic photosynthetic bacteria. Community 2 mainly consists of aerobic chemoautotrophic bacteria such as *Rhodothermaceae* from the phylum *Bacteroidetes* and *Trueperaceae* from the phylum *Thermi*, as well as anaerobic chemoautotrophic bacteria like *Anaerolineae* from the phylum *Chloroflexi*. Notably, *Acidimicrobiales* from the phylum *Actinobacteria* suggest a potential role in supporting organic compound provision through autotrophic iron ion oxidation. Additionally, *Comamonadaceae* from the phylum *Proteobacteria* is a notably abundant species. Finally, Community 3 is primarily composed of aerobic chemoautotrophic bacteria like *Arthrobacter, Nocardioidaceae*, and *Agromyces* from the phylum *Actinobacteria*. Especially, *Arthrobacter* has been reported to survive under stressful conditions (Mongodin et al., 2006). (Black, J. G., Microbiology).

## Conclusion

In our research, we crafted innovative enclosures dubbed “Ecosphere” and “Biosealed” to delve into the creation and upkeep of compact, naturally sustainable ecosystems within confined spaces. Our experiments furnished pivotal insights relevant to life support systems in space exploration:

- **Natural Circulation in Sealed Habitats:** We discerned the symbiotic synergy between plants and Cyanobacteria in the Ecosphere, which facilitate oxygen generation during daylight. This interdependence is integral to life preservation in sealed settings. Additionally, leveraging LED lighting augments plant growth, even sans sunlight—a revelation with profound implications for space missions with limited solar access.
- **Decoding Growth Dynamics:** We observed that a diminished plant growth rate correlated with ethylene gas buildup and moisture deficits. To address these challenges, an Ecosphere blueprint featuring a subterranean water reservoir has been proposed to ensure consistent moisture availability. Furthermore, variations in leaf dimensions, modulated by plant hormones, emerged as vital indicators of plant health and vitality.
- **Navigating Planetary Soil Terrains:** Under severe soil conditions, the incorporation of microorganisms seems promising in enhancing seed sprouting. This underscores the merit of microbial symbiosis as a viable strategy for life support in inhospitable extraterrestrial conditions. The soil’s intrinsic properties, such as granular size, hydrophobicity, and cohesiveness, significantly impact plant development. Recognizing these factors is vital in refining life support system architectures.
- **Microbial Dynamics in Extraterrestrial Soils:** Understanding microbial populations thriving in simulated alien soils is essential, as it could foreshadow potential pathogen proliferation during extraterrestrial colonization. Given the pivotal link between gut microbiome and factors like healthspan, this exploration is of utmost importance.

While our study provided substantial insights, there are limitations to consider. Firstly, the “Ecosphere” and “Biosealed” systems, despite emulating extraterrestrial conditions, were still influenced by Earth’s specific gravitational and atmospheric conditions. This may not entirely encapsulate the challenges posed by actual space environments. The dependence on LED lighting, while demonstrating potential, raises questions about energy requirements and the feasibility of maintaining such systems during extended space voyages. Moreover, while we have charted the behavior of specific microbial communities in our simulated environments, the full spectrum of microbial interactions and potential risks in genuine extraterrestrial contexts remains uncertain. Collectively, these findings provide a robust framework to enhance the design and functionality of life support systems in the evolving realm of space exploration.

Collectively, these findings provide a robust framework to enhance the design and functionality of life support systems in the evolving realm of space exploration.

## Materials and methods

### Preparation of the Ecosphere Container

The Ecosphere utilized in this research was crafted from a sealed glass container. Plastic containers were not chosen due to their vulnerability to heat, making them unsuitable for prolonged experiments. The lid of the container was fortified with silicone and further sealed at the mouth with Parafilm for added protection.

### Construction of Ecosphere1

Soil samples were taken from various locations in the Kumayama mountains in Okayama, Japan, and mixed with soil where clover grew. Water collected from our immediate environment was used. The collected soil, water, and plant seeds were placed in the glass container and sealed. No sterilization process was implemented.

### Construction of Ecosphere2

Large grain Akadama soil (Sup Fig. 6b) was laid at the bottom of the container. Cultivated soil (Sup Fig 6a), leaf mold, chemical fertilizer with a nitrogen (N), phosphorus (P), and potassium (K) ratio of 6:6:6, and soil from clover-growing regions were mixed and placed on top. Water from the surrounding environment was poured over the Akadama soil, followed by the mixed soil. Clover seeds were then spread over this layer and the container was sealed. The interior was composed of air, soil, and groundwater layers with respective ratios of 13:5:4.

### Construction of Biosealed

The same cultivated soil, leaf mold, and fertilizer with a ratio of 6:6:6 N:P:K, as used in Ecosphere2, were combined. To enhance moisture retention, zeolite (Sup Fig. 6c) and vermiculite (Sup Fig. 6d) were added. Eggs of the yellow Drosophila fly, cabbage, and clover seeds were introduced, and the glass container was sealed.

### Creation of Exoplanetary Simulated Soils

#### Lunar Simulated Soil

We employed a lunar simulated soil with the same composition as lunar regolith (provided by Hideaki Miyamoto, Sup Fig.5a). Refer to Sup Fig. 8 for the chemical composition.

#### Ryugu Simulated Soil

CI chondrite, possessing a similar composition to Ryugu soil, was used as the Ryugu simulated soil (provided by Hideaki Miyamoto, Sup Fig.5b). Refer to reference 13 for more information (Miyamoto et al., 2006, Yokoyama et al., 2023, Nakamura et al., 2022, Nakamura et al., 2023).

#### Lunar Regolith Simulated Soil

Fly ash and Clinker ash, by-products from coal-fired power plants resembling lunar regolith properties, were utilized (provided by Chugoku Electric Power Co.). For the chemical composition, see Sup Fig. 8 and Sup table 3.

#### Creation of Porous Fly Ash using Bentonite as a Binder

10g of Fly ash (produced by Chugoku Electric Power - Misumi Power Plant) was mixed with 1g of bentonite (Kunigel V1 from Kunimine Industries). This mixture was kneaded with 5mL of water and molded. The firing process involved a gradual increase from room temperature to 900°C over 1 hour, maintaining 900°C for 2 hours, and then naturally cooling inside the furnace, as detailed in the time program (Koizumi et. al., 1998, Sup Fig. 9).

#### Metabolome Measurement & Analysis

##### DNA Extraction from Soil Samples

Soil samples were lyophilized with VD-250R Freeze Dryer (TAITEC) and the lyophilized samples were smashed with Multibeads Shocker (Yasui Kikai) at 1,500 rpm for 2 minutes. Lysis Solution F (Nippon Gene) was added to the smashed samples. After incubation for 10 minutes at 65 C, the samples were centrifuged at 12,000 x g for 2 minutes. The supernatants were mixed with Purification Solution (Nippon Gene) and chloroform, and centrifuged at 12,000 x g for 15 minutes. DNAs of the supernatants were purified with MPure-12 system and MPure Bacterial DNA Extraction Kit (MP Bio).

##### DNA Extraction from Water Samples

Environmental DNA and cellular fragments were concentrated from water samples using Sterivex filters (Merck Millipore, 0.22 μm). Post-filtration, Lysis Solution F (Nippon Gene) was added, and the samples were shaken at 1,500 rpm for 2 minutes using the Shake Master Neo (bms). After a 10-minute incubation at 65°C, the samples were centrifuged at 12,000 x g for a minute, and the supernatant was collected. DNA was subsequently extracted using the Lab-Aid 824s DNA Extraction kit (ZEESAN).

##### Library Creation & Sequencing

The Synergy LX (Bio Tek) and QuantiFluor dsDNA System (Promega) were employed to measure the DNA solution concentration. A library was prepared using the 2-step tailed PCR method. The Synergy H1 (Bio Tek) and QuantiFluor dsDNA System were then utilized to measure the library concentration, while the Fragment Analyzer and dsDNA 915 Reagent Kit (Agilent Technologies) confirmed the library’s quality. Sequencing was executed using the MiSeq system and MiSeq Reagent Kit v3 (Illumina) with a 2x300 bp setup.

##### 16S rRNA Data Analysis

Using the FASTX-Toolkit (ver. 0.0.14), reads that matched the used primer sequences were isolated. Primer sequences in the extracted reads were removed, and the sequences were processed using various bioinformatics tools and plugins. Lineages were inferred by comparing the obtained representative sequences with the Greengene (ver. 13_8) 97% OTU database. The phylogenetic tree was constructed using the Alignment and phylogeny plugins.

##### Microbial Community Identification

During the preprocessing phase of the analysis, OTUs derived from eukaryotic organisms, specifically “chloroplasts” and “mitochondria,” were removed from the dataset. Subsequently, we identified the bacterial community and applied BALSAMICO (BAyesian Latent Semantic Analysis of MIcrobial COmmunities) to investigate the association between the bacterial community and experimental conditions (Abe et al., 2021). BALSAMICO incorporates a non-negative matrix factorization approach that considers environmental factors, thereby modeling community structure. The number of bacterial community clusters, L=3, was determined using ten-fold cross-validation in BALSAMICO. Moreover, we designated “environmental condition” as the explanatory variable in BALSAMICO, selected “Indicator-soil” as the baseline, and incorporated “ES1,” “ES2,” and “others.”

## Acknowledgments

We express our profound gratitude to Dr. Koshiro Koizumi (Nihon University) for his invaluable contribution in the creation of porous fly ash using bentonite as a binder. Special thanks to Dr. Hideaki Miyamoto (University of Tokyo) for providing lunar and Ryugu regolith simulants, which facilitated our experimental studies in the extraterrestrial soil simulant environment. We appreciate the insightful proposal by Dr. Yuya Sakai (University of Tokyo) regarding the creation of lunar regolith simulants using Fly ash. The Chugoku Electric Power Company is gratefully acknowledged for supplying EcoPowder (Fly ash), LightSand (Clinker ash), and HiBeads solidified with cement and fly ash. The opportunity to utilize scanning electron microscopes and fluorescence microscopes was possible due to Dr. Tatsuya Ikeda (National Agriculture and Food Research Organization). We owe a debt of gratitude to Dr. Kenji Kadomatsu (Nagoya University) and Dr. Mikiko Shiomi (University of Tokyo) for providing invaluable perspectives that expanded our contemplations on the research. Special recognition goes to Dr. Haruhiko Shiomi (Keio University) for granting us the chance to delve into the relationship between microorganisms and enhanced longevity. Our understanding at the molecular biology level was deepened thanks to Dr. Hideaki Kato (University of Tokyo). Dr. Natsumi Noda (Tokyo Institute of Technology) offered invaluable insights, introducing us to the deeper realms of the search for extraterrestrial life. Through Ms. Shiori Miyamoto (Nippon Veterinary and Life Science University), we became acutely aware of the challenges surrounding space habitation and gender issues, which we believe can also be applicable to confined space confrontations. This has been extremely enlightening. Thanks to Dr. Yoshiyuki Murata (Okayama University) for broadening our knowledge about ethylene treatment. The tie between space and agriculture became clearer through the insights shared by Dr. Ayako Fukunaga (National Agriculture and Food Research Organization). Thanks to Dr. Makoto Shinohara (National Agriculture and Food Research Organization) and Mr. Takuma Ishibashi (University of Tokyo), our understanding of microorganisms and space agriculture, as well as the future of space medicine, was greatly enhanced. Endless appreciation goes to Mr. Daisuke Takeda, Mr. Shuichi Gora, Mr. Tomoki Morita, and Mr. Kazuhiro Takeuchi, science teachers from Okayama Hakuryo High School, whose relentless support enabled our five-year research journey. We extend our deepest gratitude to everyone involved in this research endeavor.

## Authors contributions

T. Sato designed and preformed research. T. Sato, K. A., J. K., T. A. Y. A. Y., M. T., K. K., M. S., J. Y., S. Y., and T. Shimamura analyzed and interpreted data. T. Sato and T. Shimamura wrote the paper.

## Contact For Reagent and Resource Sharing

Further information and requests for resources and reagents should be directed to, and will be fulfilled by Tsubasa Sato and Teppei Shimamura.

