## Supplementary Figures and Tables for "Survivability and Life Support in Sealed Mini-Ecosystems with Simulated Planetary Soils"

### Supplemental Inventory

**Supplementary Figures and Tables**

**Supplementary Figure 1**
**Supplementary Figure 2**
**Supplementary Figure 3**
**Supplementary Figure 4**
**Supplementary Figure 5**
**Supplementary Figure 6**
**Supplementary Figure 7**
**Supplementary Figure 8**
**Supplementary Figure 9**
**Supplementary Table 1**
**Supplementary Table 2**
**Supplementary Table 3**

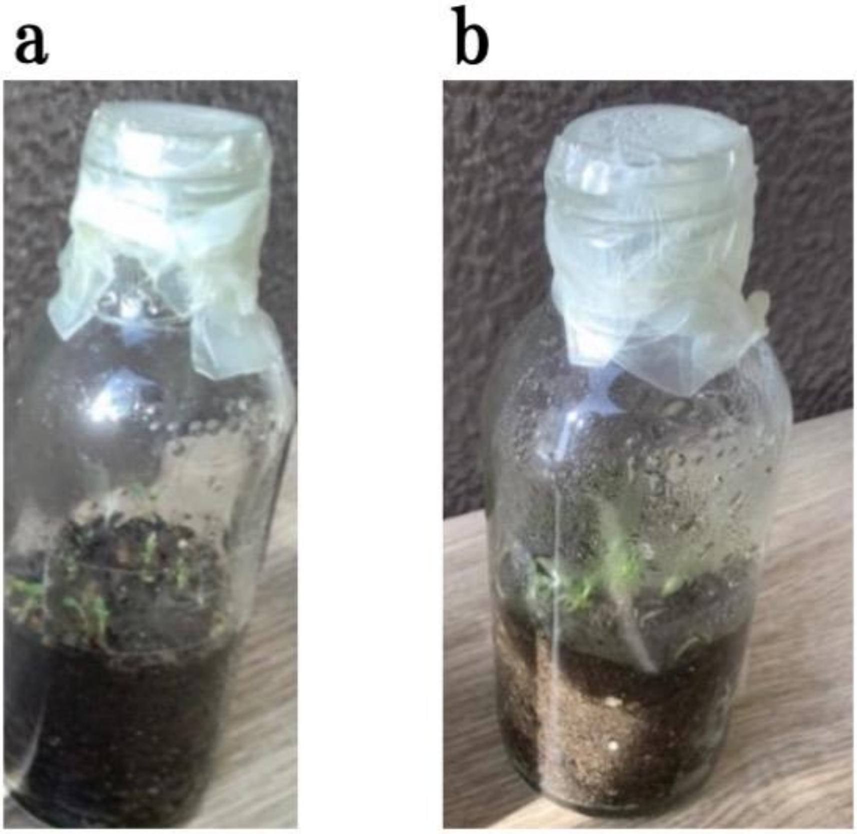


**Supplementary Figure 1: Appearance of Ecosphere 1 Used in the Mass Conservation Experiment.**

(a) Exterior of Ecosphere 1 (Sample 1) used for the mass conservation experiment. (b) Exterior of Ecosphere 1 (Sample 2) used for the mass conservation experiment.

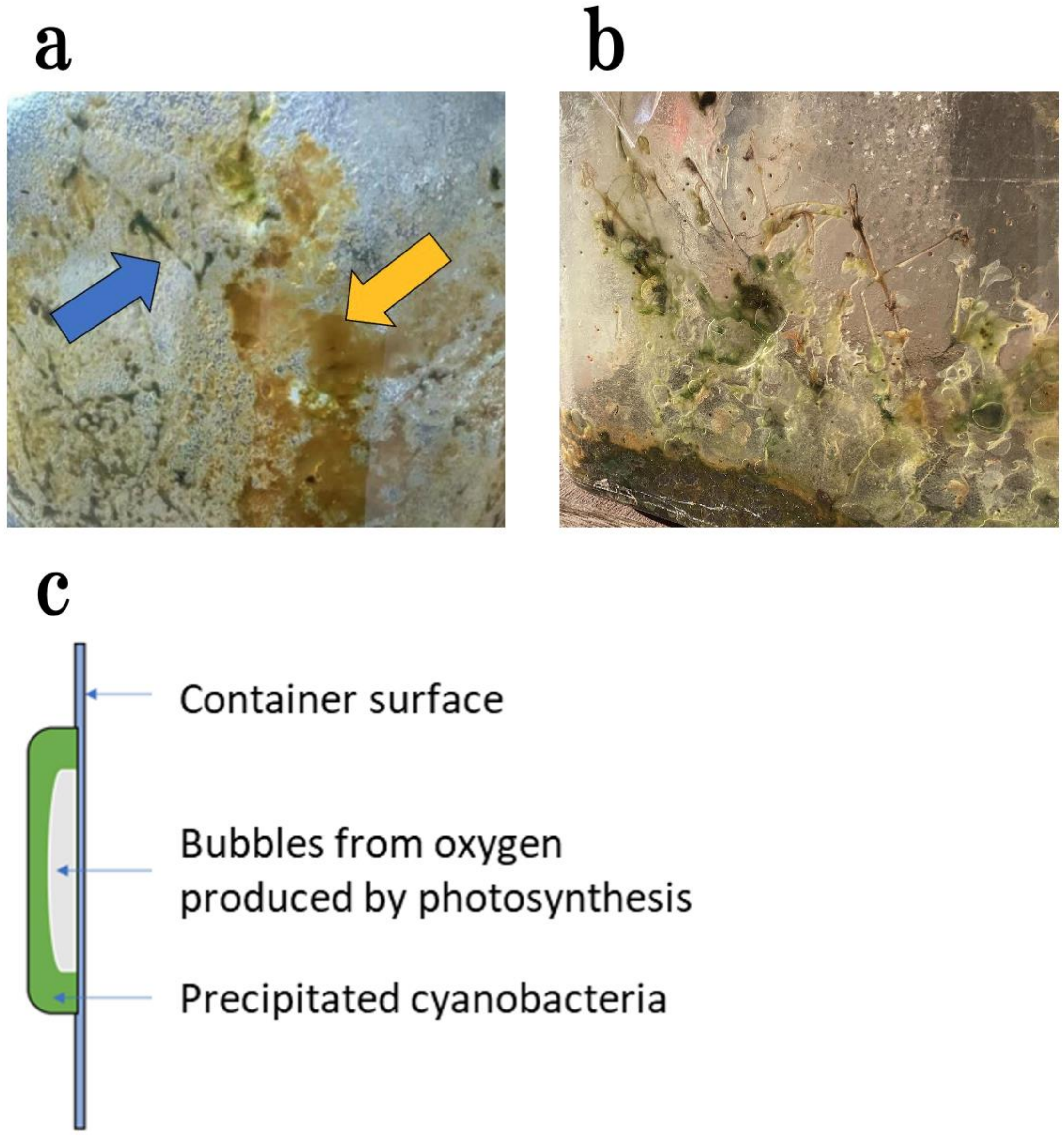


**Supplementary Figure 2: Observable Microorganisms in Ecosphere 1**

(a) Microorganisms proliferating within Ecosphere 1, visibly observed (Blue arrow indicates decomposition of stems, yellow arrow indicates the proliferation of jellyfish-like organisms). (b) Oxygen bubbles produced by cyanobacteria through photosynthesis. (c) Diagram showing that the bubbles on the container surface were generated by the photosynthesis of cyanobacteria.

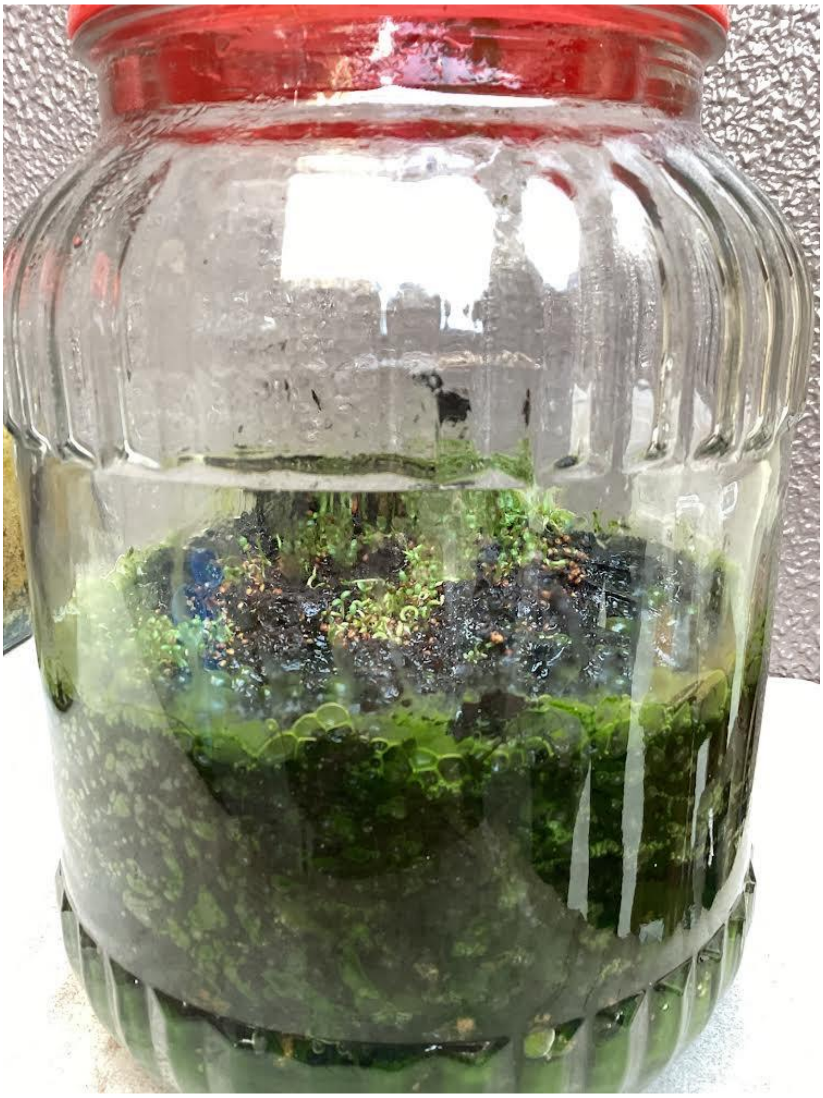

**Supplementary Figure 3: Appearance of Ecosphere 2**

With a groundwater layer created using red clay at the bottom.

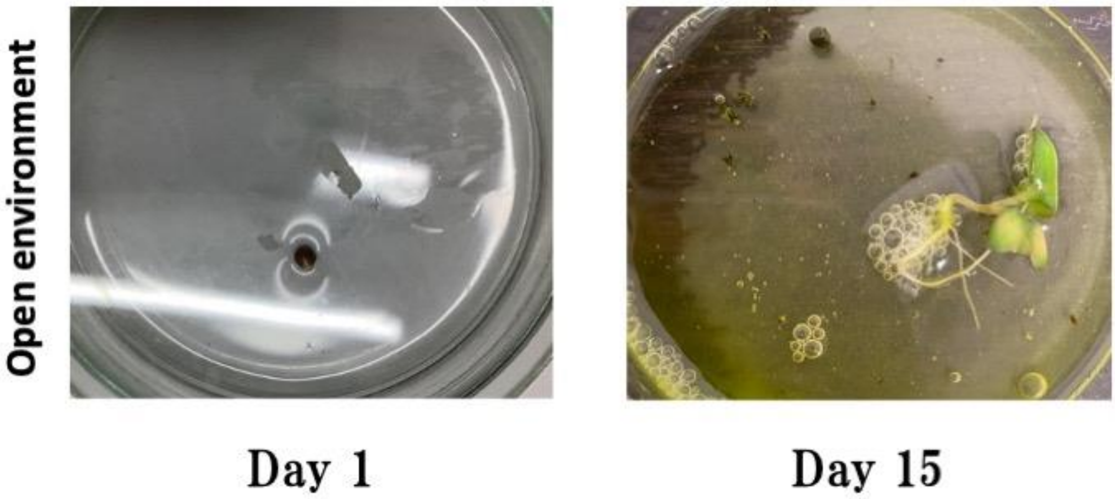


**Supplementary Figure 4: Plant Growth Under LED Light Conditions**

Observations of plant growth experiment using LED light in an open environment without soil, utilizing cabbage.

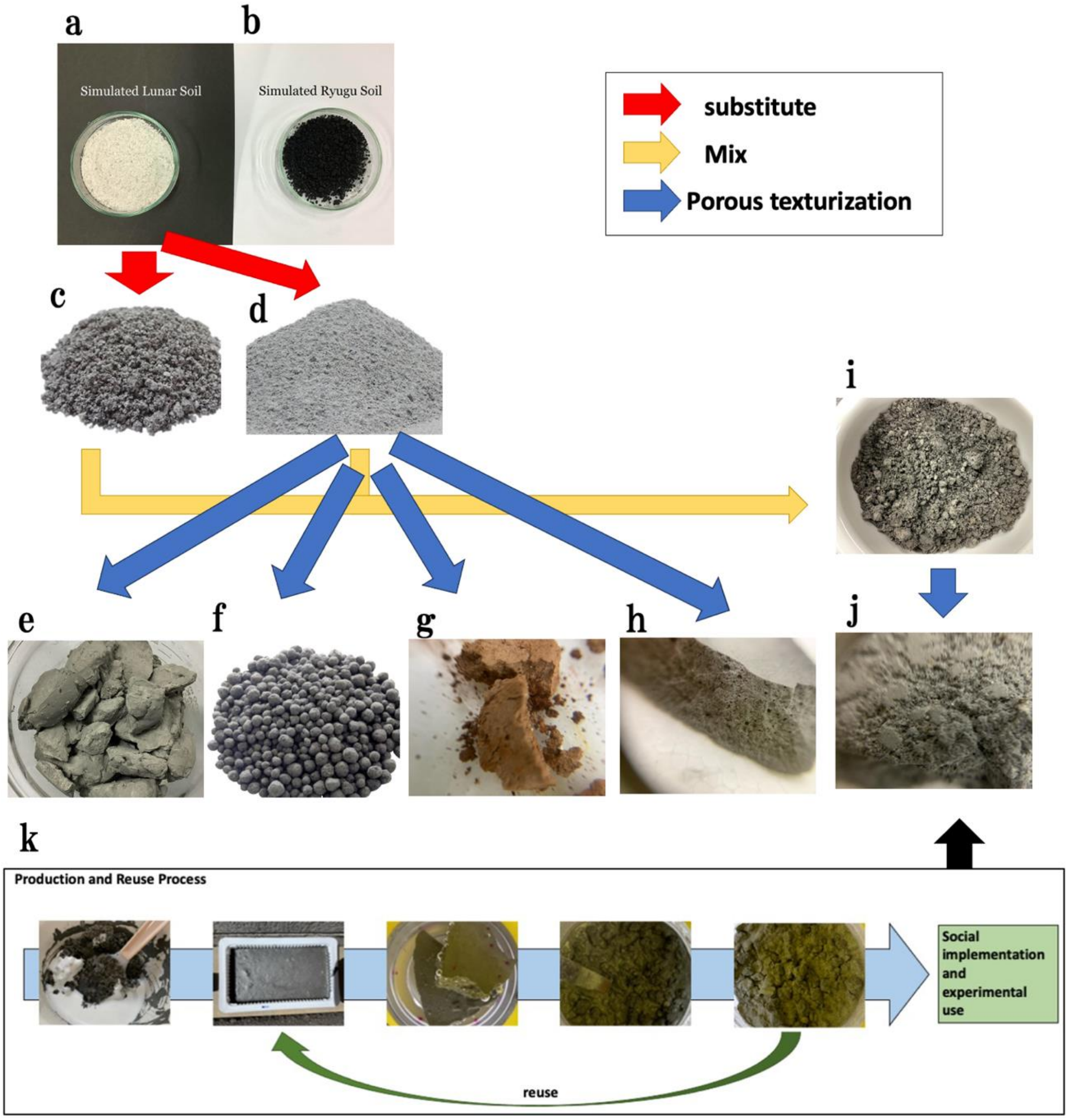


**Supplementary Figure 5: Overview of Simulated Extraterrestrial Soil and Its Modification Methods Used in the Experiment:**

(a) Simulated lunar soil (similar composition). (b) Simulated Ryugu soil. (c-d) Types of coal ash similar in characteristics and composition to lunar regolith: Clinker ash and Fly ash. (e) Lump of fly ash mixed with water. (f-h) Various modifications of fly ash using different binding agents. (i-j) Mixed soil of fly ash and clinker ash, including a porous version using a bonding agent. (k) Diagram showing the creation stages and recycling process of Boncrete.

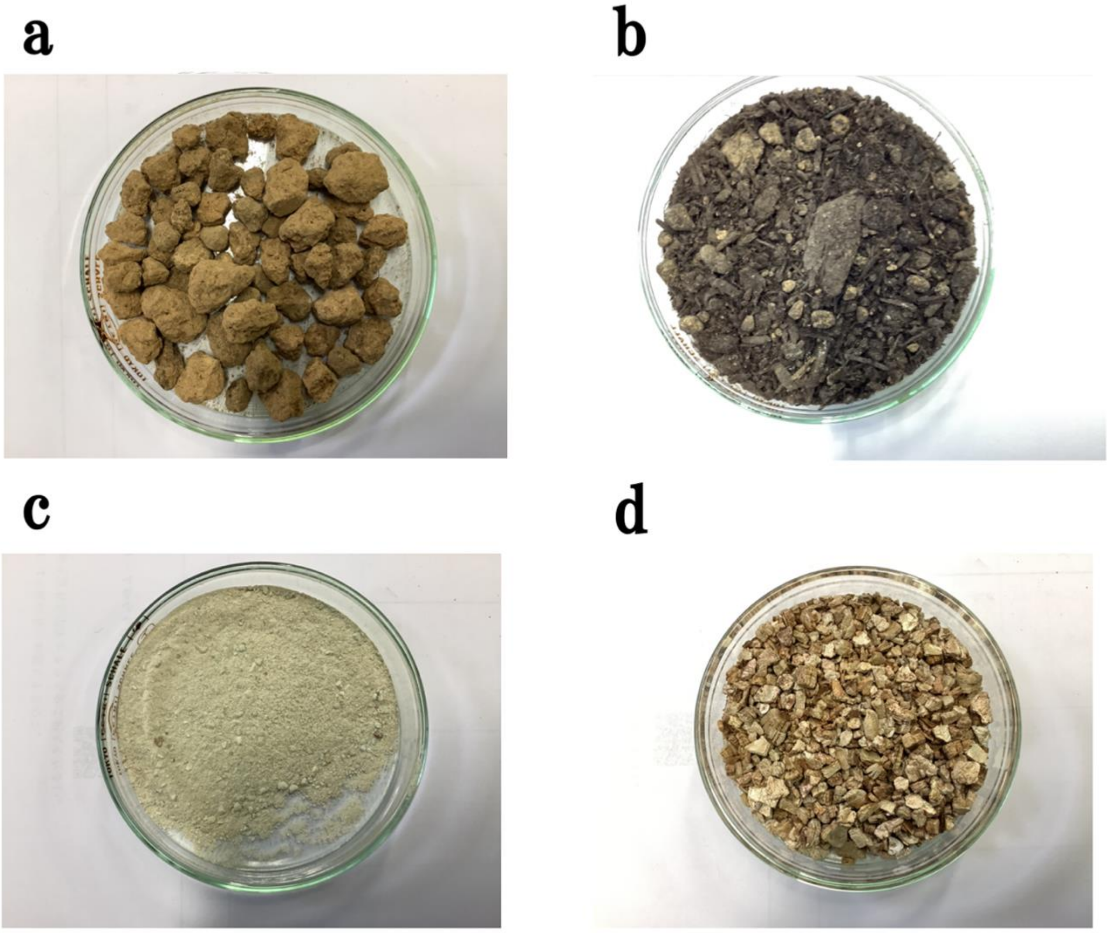


**Supplementary Figure 6: Soil Used in Ecosphere 2**

(a-b) Appearance of the soil used in Ecosphere 2. (c) Zeolite used in Ecosphere 2. (d) Vermiculite used in Ecosphere 2.

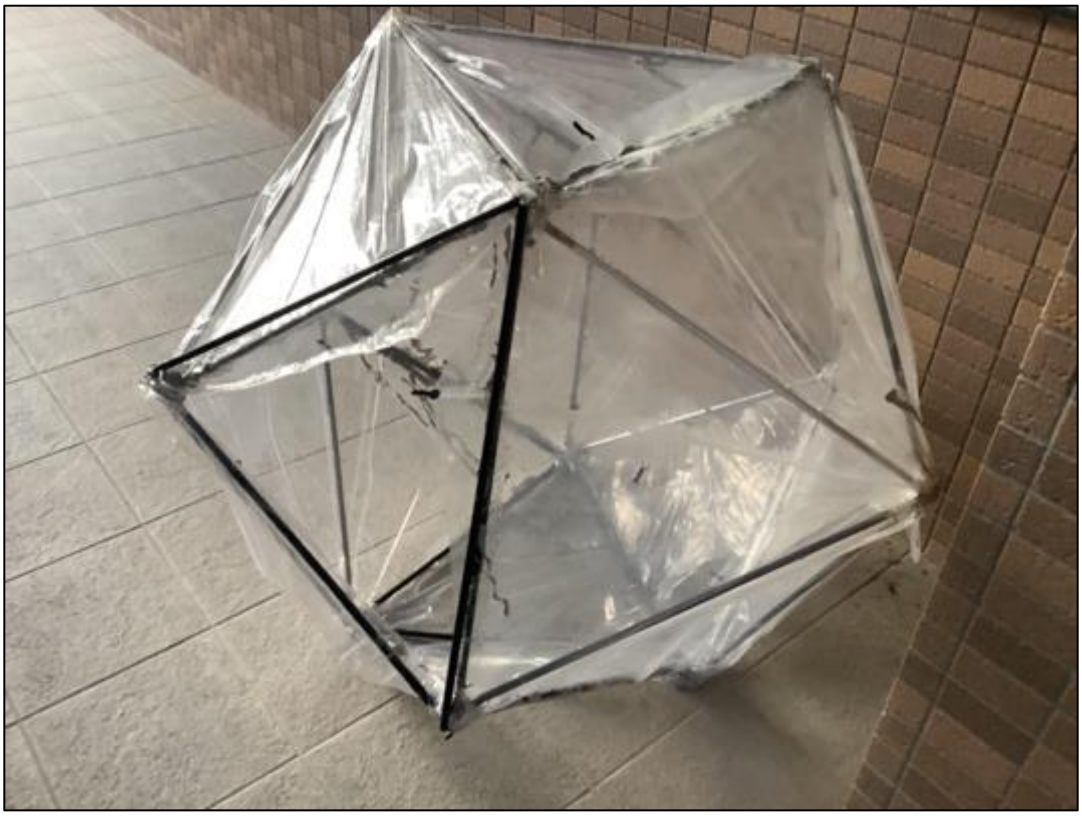

**Supplementary Figure 7: Development of Biosealed 2**

A larger version of Ecosphere and Biosealed designed to prevent entropy collapse in a small enclosed space. The structure and vinyl from an umbrella were separated, skeleton cut to 60cm sides, reinforced with string, and vinyl was secured around using silicone.

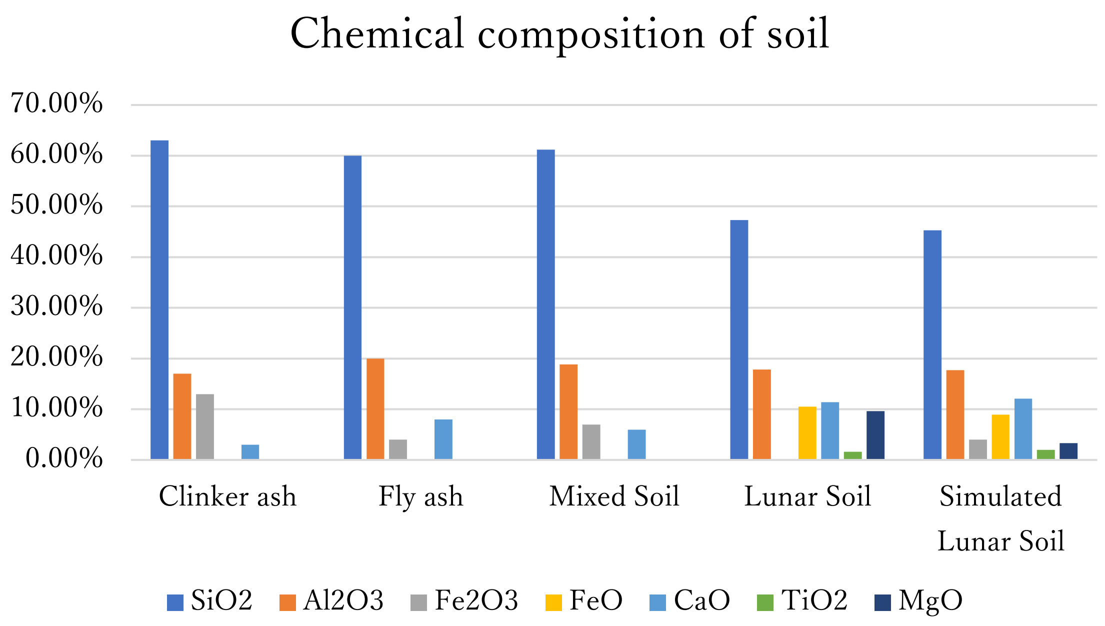


**Supplementary Figure 8: Chemical Composition of Various Soils**

Bar graph of the chemical composition of fly ash, clinker ash, their mixed soil, lunar soil, and simulated lunar soil.

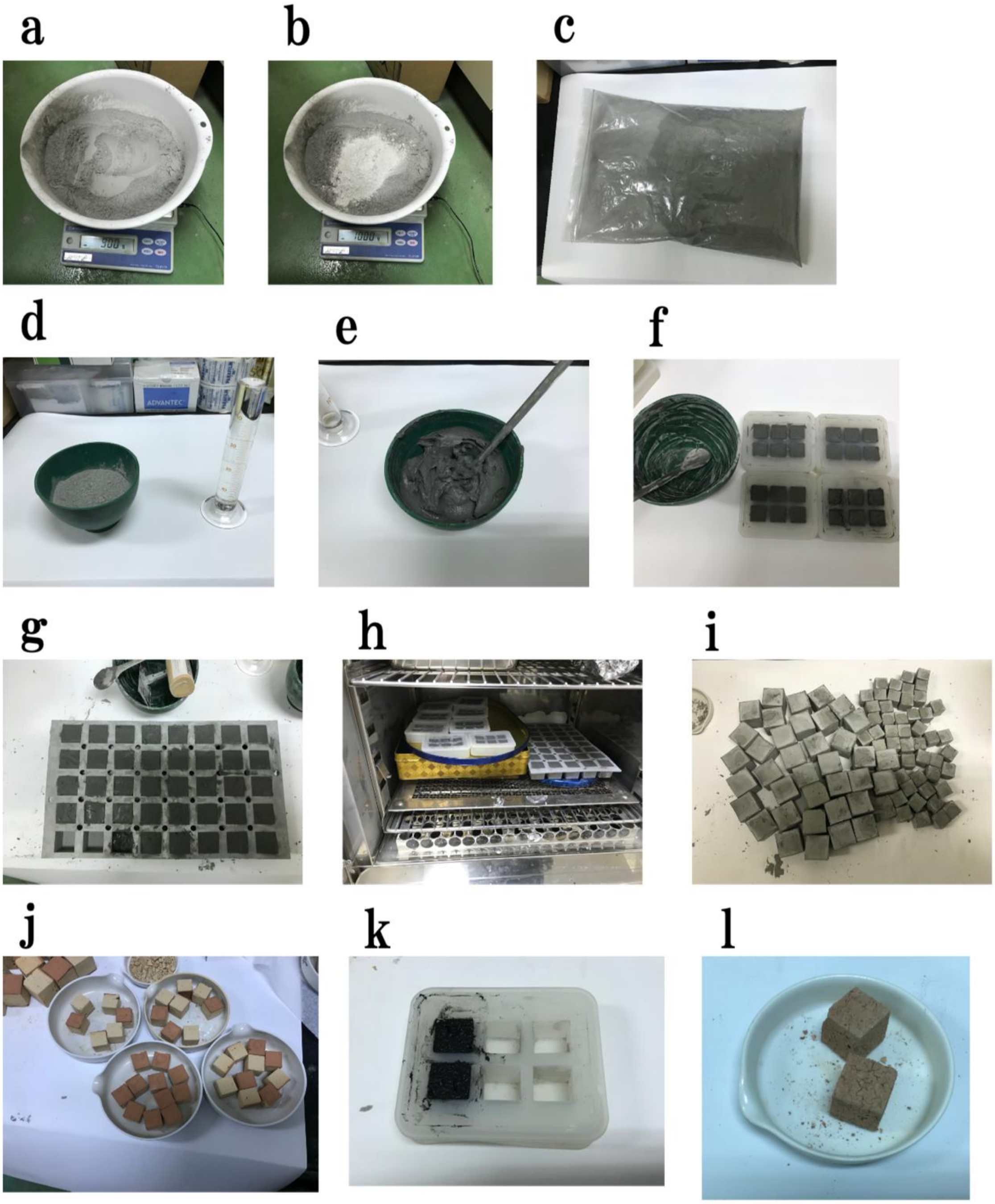

**Supplementary Figure 9: Creation of Porous Fly Ash Using Bentonite as a Binder**

(a-l) Various steps in the process, from sampling to drying and firing.

**Supplementary Table 1: Results of the Mass Conservation Experiment**

| | Before (2021/11/27) | After (2022/4/28) |
|---|---|---|
| **Sample 1** | 365g | 365g |
| **Sample 2** | 431g | 431g |

**Supplementary Table 2: Basic Information of Ecosphere 1 Used in the Mass Conservation Experiment**

| Sample Type | Sample 1 | Sample 2 |
|---|---|---|
| **Number of seeds** | 10 | 20 |
| **Water content** | 20 ml | 20 ml |
| **Soil capacity** | 50 ml | 50 ml |
| **Air capacity** | 65 ml | 65 ml |
| **Total capacity** | 115 ml | 115 ml |

**Supplementary Table 3: Chemical Composition of Fly ash, Clinker ash, their mixed soil, lunar soil, and simulated lunar soil**

| | Clinker ash | Fly ash | Mixed Soil | Lunar Soil | Simulated Lunar Soil |
|---|---|---|---|---|---|
| $SiO_2$ | 63.00% | 60.00% | 61.17% | 47.30% | 45.30% |
| $Al_2O_3$ | 17.00% | 20.00% | 18.82% | 17.80% | 17.70% |
| $Fe_2O_3$ | 13.00% | 4.00% | 7.00% | — | 4.00% |
| FeO | — | — | — | 10.50% | 8.90% |
| CaO | 3.00% | 8.00% | 5.96% | 11.40% | 12.10% |
| $TiO_2$ | — | — | — | 1.60% | 2.00% |
| MgO | — | — | — | 9.60% | 3.30% |
